# Frontal-sensory cortical projections become dispensable for attentional performance upon a reduction of task demand in mice

**DOI:** 10.1101/2021.04.08.439093

**Authors:** Kevin J. Norman, Julia Bateh, Priscilla Maccario, Christina Cho, Keaven Caro, Tadaaki Nishioka, Hiroyuki Koike, Hirofumi Morishita

## Abstract

Top-down attention is a dynamic cognitive process that facilitates the detection of the task-relevant stimuli from our complex sensory environment. A neural mechanism capable of deployment under specific task-demand conditions would be crucial to efficiently control attentional processes and improve promote goal-directed attention performance during fluctuating attentional demand. Previous studies have shown that frontal top-down neurons projecting from the anterior cingulate area (ACA) to the visual cortex (VIS; ACA_VIS_) are required for visual attentional behavior during the 5-choice serial reaction time task (5CSRTT) in mice. However, it is unknown whether the contribution of these projecting neurons is dependent on the extent of task demand. Here, we first examined how behavior outcomes depend on the number of locations for mice to pay attention and touch for successful performance, and found that the 2-choice serial reaction time task (2CSRTT) is less task demanding than the 5CSRTT. We then employed optogenetics to demonstrate that suppression ACA_VIS_ projections immediately before stimulus presentation has no effect during the 2CSRTT in contrast to the impaired performance during the 5CSRTT. These results suggest that ACA_VIS_ projections are necessary when task demand is high, but once a task demand is lowered, ACA_VIS_ neuron activity becomes dispensable to adjust attentional performance. These findings support a model that the frontal-sensory ACA_VIS_ projection regulates visual attention behavior during specific high task demand conditions, pointing to a flexible circuit-based mechanism for promoting attentional behavior.

## 1. Introduction

Top-down attention is a fundamental cognitive process that facilitates the detection of the most pivotal goal-directed stimuli from our dynamic environment. The frontal cortex, particularly the anterior cingulate cortex area (ACA), has been demonstrated to be a key mediator of top-down control of visual attention across species including rodents (Passetti et al., 2002; Chudasama et al., 2003; Pehrson et al., 2013; Kim et al., 2016; Koike et al., 2016). Among ACA neurons, recent studies in mice have demonstrated that frontal-sensory projections from the ACA to the primary visual cortex (VIS) (ACA_VIS_) modulate visual discrimination (Zhang et al., 2014; Moore and Zirnsak, 2017) and free-moving attentional behavior (Norman et al., 2021a) through modulation of visual cortex processing (Norman et al., 2021b). However, it is not known whether the contribution of this top-down circuit depends on task demand. A circuit mechanism capable of deployment under specific task-demand conditions would be crucial to efficiently control attentional processes and improve promote goal-directed attention performance during fluctuating attentional demand. Here, we aimed to determine how task demand impacts the contribution of frontal-sensory projection neurons to attentional behavior.

We hypothesized that ACA_VIS_ projection neurons may only be necessary for visual attentional behavior under conditions of elevated task demand. We tested this hypothesis by subjecting mice to two tasks of varying task demand while selectively manipulating ACA_VIS_ neural activity in mice. One way to impact the demand of anticipatory attention is to vary the number of locations for mice to scan and touch for successful performance. Here we used the tasks in which mice were required to sustain and divide their attention in anticipation of a random presentation of a brief stimulus at one location across select number of response windows; either 5 locations during the 5CSRTT (Carli et al., 1983) (5CSRTT, **Fig. S1**) or limited to 2 locations during the 2CSRTT (Dillon et al., 2009; van Gaalen et al., 2009). During these tasks, we employed optogenetics to suppress ACA_VIS_ projection activity during these tasks to assess to what extent task demand impacts the contribution of ACA_VIS_ projection neurons to visual attentional behavior.

## 2. Materials and Methods

### 2.1 Experimental model

Adult, male C57Bl/6 mice (Charles River Laboratories, MA) were group-housed under a standard 12 hr light/dark cycle in a temperature- and humidity-controlled vivarium. Training was initiated when mice were 9–10 weeks old. Mice were allowed access to water for 2 hours each day and maintained approximately 85-90% of their *ad libitum* weight during behavioral training. Food was available *ad libitum* throughout the experiment. All animal protocols were approved by the Institutional Animal Care and Use Committee at Icahn School of Medicine at Mount Sinai. Data of the 5CSRTT have been analyzed previously (Norman et al., 2021b), but only the mice which underwent both 5CSRTT and 2CSRTT with optogenetics (6 out of 8 mice) were re-analyzed in this study.

### 2.2 Viral Strategies and Stereotaxic Procedures

Following procedures previously described in (Norman et al., 2021b), mice were anesthetized with 2% isoflurane and head-fixed in a mouse stereotaxic apparatus (Narishige, East Meadow, NY). AAV2-CamKII-eNpHR3.0-eYFP (UNC Viral Vector Core, Chapel Hill, NC) was injected bilaterally into the ACA. Bilateral ACA injection sites relative to Bregma area are: AP +0.7mm, ML ±0.2mm, DV −0.7mm; AP +0.2mm, ML ±0.2mm, DV −0.7mm; AP −0.3mm, ML ±0.2mm, DV −0.7mm. Bilateral VIS injection sites relative to lambda are: AP +0.0mm, ML ±3.0mm, DV −0.4mm; AP +0.1mm, ML ±2.85mm, DV −0.4mm; AP +0.1mm, ML ±3.15mm, DV −0.4mm. Each infusion (500 nl) was made at 150 nl/min using a microinjector set (Nanoject III) and glass pulled syringe. The glass pipettes (1.14 mm outer diameter and 0.53 mm inner diameter, 3-000-203-G/X, Drummond Scientific, PA) were pulled on a P-97 Flaming/Brown type micropipette puller (Sutter Instrument, CA). The tip of the pulled glass pipettes was approximately 50 μm. The syringe was left in place for 1min following the injection to reduce backflow of virus. Bilateral LEDs (Amuza, San Diego, CA) of 500μm diameter that delivered 470nm light were implanted at the VIS. A glass cannula associated with LED was implanted at the region of interest, and was glued onto the brain and light from the LED traveled about 1mm through glass cannula into the brain. Model info of the LED and light intensity info is also included. Behavioral testing occurred at least three weeks after viral injection to allow for maximal viral expression. The location of the cannula was mapped onto blank coronal slice templates taken from Paxinos and Franklin mouse brain atlas as previously described (Norman et al., 2021b). One mouse was excluded due to the off-target cannula implantation.

### 2.3 Behavior

#### 5- and 2-Choice Serial Reaction Time Task (5CSRTT/2CSRTT)

##### Apparatus

5CSRTT and 2CSRTT (See also **Fig. S1**) was conducted in eight Bussey–Saksida operant chambers with a touchscreen system (Lafayette Instruments, Lafayette, IL) following procedures previously described in (Norman et al., 2021b). Dimensions are as follows: a black plastic trapezoid (walls 20 cm high × 18 cm wide (at screen-magazine) × 24 cm wide (at screen) × 6 cm wide (at magazine)). Stimuli were displayed on a touch-sensitive screen (12.1 inch, screen resolution 600 × 800) divided into five response windows by a black plastic mask (4.0 × 4.0 cm, positioned centrally with windows spaced 1.0 cm apart, 1.5 cm above the floor) fitted in front of the touchscreen. For the 5CSRTT, all five response windows were available, but during the 2CSRTT, three of the outer response windows were masked and only two adjacent center windows were available. Schedules were designed and data was collected and analyzed using ABET II Touch software (v18.04.17, Lafayette Instrument). The inputs and outputs of the multiple chambers were controlled by WhiskerServer software (v4.7.7, Lafayette Instrument).

##### Habituation

Following procedures previously described in (Norman et al., 2021b), before 5CSRTT training, mice were first acclimated to the operant chamber and milk reward. The food magazine was illuminated and diluted (30%) sweetened condensed milk (Eagle Brand, Borden, Richfield, OH) was dispensed every 40s after mice entered the food magazine. Mice needed to enter the reward tray 20 times during two consecutive 30min sessions before advancing to the next stage. Mice were then trained to touch the illuminated response window. During this phase, a white square stimulus was presented randomly at one response window until it was touched. If the mouse touched the stimulus, the milk reward was delivered in conjunction with a tone and magazine light. Touches to nonstimuli had no consequence. After reaching criterion on this phase (20 stimulus touches in 30 min for 2 consecutive days), mice advanced to the 5CSRTT training phase.

##### 5CSRTT Training and Baseline

Following procedures previously described in (Norman et al., 2021b), mice were tested 5 days a week, 100 trials a day (or up to 30min). Each trial began with the illumination of the magazine light. After mice exited the food magazine, there was an intertrial interval (ITI) period of 5s before a stimulus was presented randomly at one response window. If a mouse touched the screen during the ITI period, the response was recorded as premature and the mouse was punished with a 5s time-out (house light on). After the time-out period, the magazine light illumination and house light switch off signaled onset of the next trial. After the ITI period, a stimulus appeared randomly in one of the five response windows for a set stimulus duration (this varied from 32 to 2s, depending on stage of training). A limited-hold period followed by the stimulus duration was 5s, during which the stimulus was absent but the mouse was still able to respond to the location. Responses during stimulus presence and limited holding period could be recorded either as correct (touching the stimulus window) or incorrect (touching any other windows). A correct response was rewarded with a tone, and milk delivery, indicated by the illumination of the magazine light. Failure to respond to any window over the stimulus and limited-hold period was counted as an omission. Incorrect responses and omissions were punished with a 5-s time-out. In addition, repeated screen touches after a correct or incorrect response were counted as perseverative responses. Animals started at stimulus duration of 32s. With a goal to baseline mice at a stimulus duration of 2s, the stimulus duration was sequentially reduced from 32, 16, 8, 4, to 2s. Animals had to reach a criterion (≥50 trials, ≥80% accuracy, ≤20% omissions) over 2 consecutive days to pass from one stage to the next. After reaching baseline criterion with the 2s stimulus duration for 4 out of 5 days, mice began 5CSRTT testing.

##### 5CSRTT Testing

Following procedures previously described in (Norman et al., 2021b), during 5CSRTT testing, attention demand was increased by reducing and pseudo-randomly shuffling the stimulus duration to 2.0, 1.5, 1.0, and 0.8s for 4 out of 5 days. Following 5CSRTT testing, mice underwent stereotaxic viral surgery and then were reestablished to baseline criterion before optogenetic experiments (2.0 and 1.0s stimulus duration). Between experimental testing days, mice were subjected to 2s stimulus duration training to confirm that the mice maintain stable baseline criterion. Attention and response control were assessed by measuring the following performance: correct percentage ((100 x (correct responses)/(correct responses + incorrect responses + omissions)), percentage accuracy (100 × correct responses/(correct responses + incorrect responses)), percentage omission (100 × omissions/(omissions + correct responses + incorrect responses)), percentage of premature responses, percentage of perseverative responses, latency to correct response (s), and latency to reward collection (s) after correct choices. A premature response is a response given during the ITI before a stimulus appears. It is a measure of impulsive behavior. A perseverative response is when a response to any screen continues to be given after the correct or incorrect screen has been poked and before retrieval of the reward. It is a measure of compulsive behavior. Correct response latency (CRL) is the difference between the time of stimulus onset and the time of the correct response. It is a measure of attention and processing speed. Reward collection latency (RCL) is the difference between the time of a correct response and the time of reward collection and reflects motivation.

##### 2CSRTT Baseline and Testing

Immediately following 5CSRTT testing, the mice underwent baselining for the 2CSRTT. During the 2CSRTT, 3 of the response windows were blocked and only 2 central response windows remained in which the stimulus could appear. Animals had to reach a baseline criterion (≥50 trials, ≥80% accuracy, ≤20% omissions) with the 2s stimulus duration at 1 of 2 locations for 4 out of 5 days to pass criteria before beginning 2CSRTT testing. During 2CSRTT testing, attention demand was increased by reducing and pseudo-randomly shuffling the stimulus duration to 2.0, 1.5, 1.0, and 0.8s for 2 consecutive days. A separate cohort of naïve mice underwent 2CSRTT training and testing, without prior 5CSRTT experience, using the same criteria as the 5CSRTT but with only 2 central response windows.

### 2.4 Optogenetics

Following procedures previously described in (Norman et al., 2021b), optogenetic experiments were conducted using Teleopto system (Amuza, San Diego, CA). Transistor-transistor logic signals were sent to a signal generator (Rigol Technologies, Beijing, China) that drove the light at specific time points during the trials. During separate test sessions, bilateral blue LED cannula with1mm length of optic fiber and 1.3mm for the center to center distance of optic fibers (TeleLCD-B-1000-500-1.3, Amuza, San Diego, CA) were stimulated during either −5:−2s of ITI, −3:0s of ITI or during the length of the stimulus presentation. One timing of stimulus duration was used per test day in a counterbalanced manner. During behavioral 5CSRTT and 2CSRTT, continuous 470nm light was delivered bilaterally via LED optic fiber implanted at the VIS pseudo randomly at specific time points throughout a trial during 50% of trials. In our study, we chose to use 470nm LED to suppress eNpHR3.0-expressing ACA neurons, because previous studies found that red-shifted light frequencies resulted in off-target effects in control mice that reduced 5CSRTT performance (White et al., 2018). During optogenetics testing, stimulus duration was pseudo randomly shuffled between 2.0 and 1.0 s. The power at the fiber-optic tip measured using an optical power meter (Thorlabs) was approximately 10 mW.

### 2.5 Quantification and Statistical Analysis

Statistical analyses were performed using Prism (Graphpad, San Diego, CA) and in R v. 3.5.3. Analyses were conducted using general linear mixed models, ANOVA, or t-tests as indicated. Bar graphs are represented as the mean and error bars represent standard error of the mean (s.e.m.)

## 3. Results

### 3.1 2CSRTT shows reduced task demand compared to 5CSRTT

We first sought to determine whether attentional behavior outcomes depend on the extent of task demand by manipulating a number of locations where visual stimulus is presented for mice to scan and touch to successfully obtain food reward. Specifically, we compared the performance between mice performing the 5CSRTT (Carli et al., 1983; Bari et al., 2008) and 2CSRTT with varying numbers of choice locations. Mice were first trained to perform the 5CSRTT (**Fig 1A, S1A**). The task required mice to sustain and divide their attention across 5 response windows in their lower visual field in anticipation of a brief white square stimulus appearing pseudorandomly at 1 of the 5 locations in automated standardized Bussey-Saksida touch-screen operant chambers (Bussey et al., 2012; Mar et al., 2013). The number of trials with correct touches to the flashed screen, compared to omitted and incorrect touches is a primary operational definition of attention. Correct response rate provides a broad and general index of attention performance and reflects what is captured by both accuracy and omission, two readouts that have been used to assess some aspects of attention in previous studies using 5CSRTT (Carli et al., 1983). It should be noted that omission probability is sensitive to both lapse in attention and motivation and every operant response depends to some degree on both. However, another readout of the 5CSRTT, the latency to collect reward following a correct response, serves as an indicator of motivation to assess if omission change reflects attention or motivation level. In addition, other 5CSRTT behavioral measures such as premature, and preservative response assess relatively independent indices of cognition (Dalley et al., 2004; Lustig et al., 2013; Koike et al., 2016) (**Fig.S1B**). Mice had to reach baseline criteria of 4 out of 5 sessions with ≥50 trials, ≥80% accuracy, ≤20% omission at 2s stimulus duration before 5CSRTT testing in which the stimulus duration length was shuffled between 2.0, 1.5, 1.0, and 0.8 s in a pseudorandomized order. Following 5CSRTT testing, mice began 2CSRTT baselining with 2s stimulus duration in which the number of response windows was reduced to 2. To pass 2CSRTT baselining stage, mice had to reach same baseline criteria as the 5CSRTT before undergoing 2CSRTT testing. We found that task performance of mice was significantly higher during the 2CSRTT compared to the 5CSRTT as indicated by increased correct trials **(Fig.1B**: two-way repeated measures analysis of variance (RM ANOVA) task type *P*=0.0059). The improved task performance during the 2CSRTT was driven by improved accuracy (**Fig.1C**: two-way RM ANOVA, task type *P*=0.0118) with a lesser degree of change in omissions (**Fig.1D**: two way RM ANOVA, task type *P*=0.1146). Furthermore, correct response latency was slower during the 5CSRTT (**Fig.1E:** t test *P*=0.0002), while reward collection latency(s) (**Fig.1F:** t test *P*=0.7720), premature responses (%) **(Fig.1G:** t test *P*=0.3218), and perseverative responses (%) (**Fig.1H**: t test *P*=0.2377) were not statistically different between both the 5CSRTT and 2CSRTT. These data suggests that each task engaged similar levels of motivation, impulsivity, and compulsivity, respectively.

**Figure 1.**
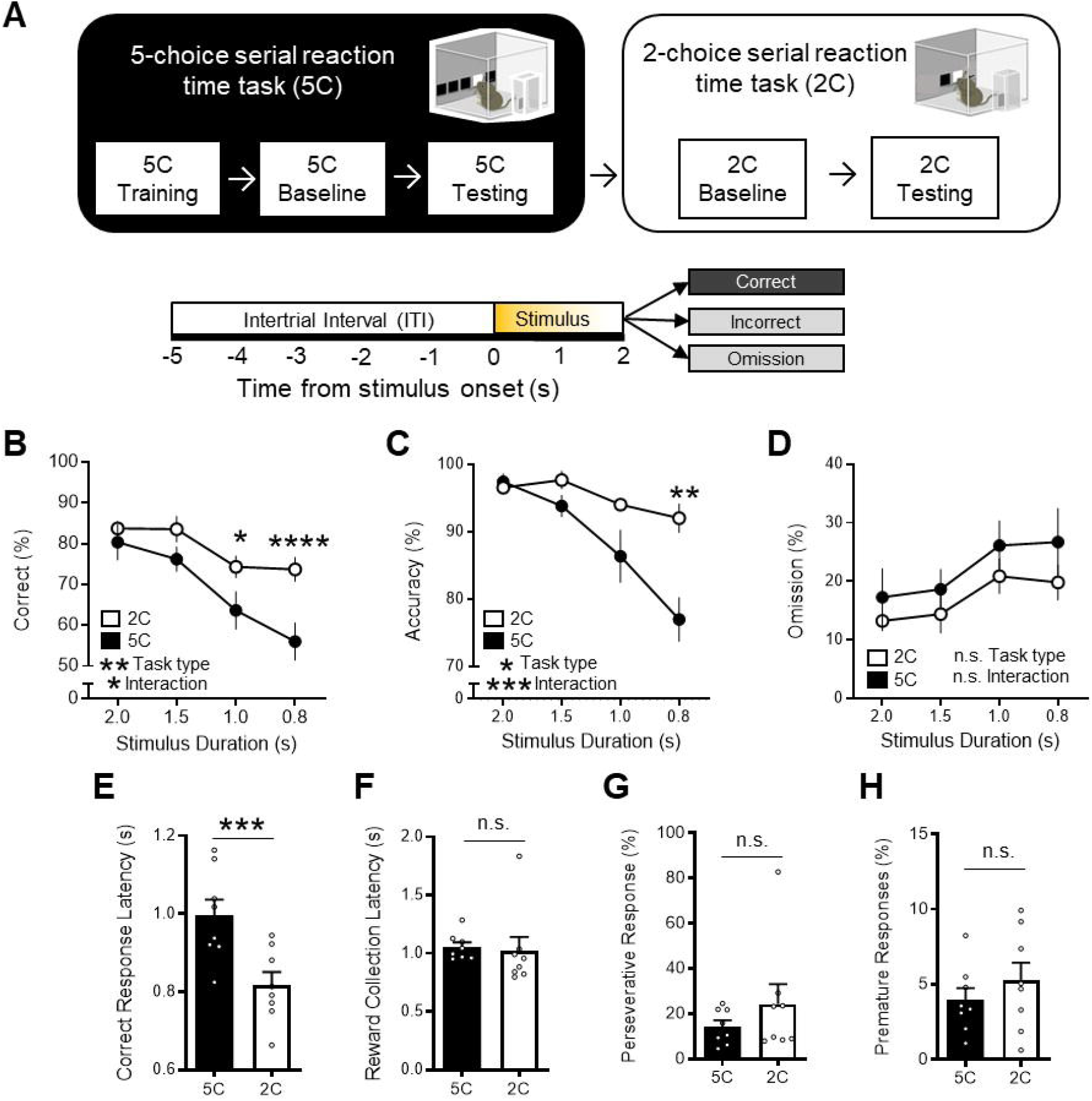
2CSRTT is less task demanding than the 5CSRTT. **(A)** Experimental timeline: Mice were first trained on the 5CSRTT before undergoing 5CSRTT testing. During 5CSRTT testing, with a fixed 5s ITI and pseudorandomized stimulus duration (2.0, 1.5, 1.0, or 0.8s), (8 mice; 2,161 total trials). The same mice were then baselined with 2CSRTT before undergoing 2CSRTT testing (2,354 total trials). **(B)** Mice had a significantly increased correct trials during the 2CSRTT compared to the 5CSRTT(%, two-way repeated measures analysis of variance (RM ANOVA), Main effect (task type) F_1,7_ =15.26, \*\**P*=0.0059, Main effect (stimulus duration): F_3,21_=29.48, *P*<0.0001, Interaction: F_3,21_ =3.626, \**P*=0.0298, Holm-Sidak multiple comparisons at 2.0, 1.5, 1.0, 0.8 s stimulus duration, *P*=0.7623, 0.1197, *0.0121, ****<0.0001, n=8 mice). **(C)** Mice had a significantly increased accuracy during the 2CSRTT compared to the 5CSRTT (%, two-way RM ANOVA, Main effect (task type): F_1,7_ =11.40, \**P*=0.0118, Main effect (stimulus duration): F_1.53,10.71_ =31.92, *P*<0.0001, Interaction: F_3,21_ =10.64, \*\*\**P*=0.0002, Holm-Sidak multiple comparisons at 2.0, 1.5, 1.0, 0.8 s stimulus duration, *P*=0.9781, 0.2574, 0.3198, **0.0017, n=8 mice). **(D)** Mice had no difference in omissions during the 2CSRTT compared to the 5CSRTT(%, two-way RM ANOVA, Main effect (task type): F_1,7_ =3.246, *P*=0.1146, Main effect (stimulus duration): F_3,21_ =9.526, *P*=0.0004 Interaction: F_3,21_ =0.2317, *P*=0.8733, n=8 mice). **(E)** The 2CSRTT had a significantly reduced correct response latency compared to the 5CSRTT (t_7_=7.193, \*\*\**P*=0.0002). **(F-H)** No difference in **(F**) reward collection latency (t_7_=0.3012, *P*=0.7720), **(G**) premature responses (t_7_=1.066, *P*=0.3218), and **(H)** perseverative responses (t_7_=1.291, *P*=0.2377) was observed between 2CSRTT and 5CSRTT testing. Error bars indicate mean□±□s.e.m. n.s. = non significant. * indicates *P*<0.05, ** indicates *P*<0.01, *** indicates *P*<0.001, **** indicates *P*<0.0001.

Of note, there was no difference in 2CSRTT performance between mice with or without prior 5CSRTT testing (**Fig. 2)**. Specifically, neither correct % (**Fig. 2B:** two-way RM ANOVA, task type *P*=0.9379), accuracy % (**Fig. 2C:** two-way RM ANOVA, task type *P*=0.9489) nor omission % (**Fig. 2D:** two-way RM ANOVA, task type *P*=0.9383) was different during 2CSRTT between mice with or without prior 5CSRTT testing. No difference in correct response latency (**Fig. 2E:** t test *P*=0.3119), reward collection latency (**Fig. 2F:** t test *P*=0.9363), premature responses (**Fig. 2G:** t test *P*=0.2913), and perseverative responses (**Fig. 2H:** *P*=0.1111) was observed between the two groups. These results suggest that the order of training has limited influence to attentional behavior. Collectively, these data support that lower task demand of 2CSRTT compared to 5CSRTT is reflected to better task performance in 2CSRTT.

**Figure 2.**
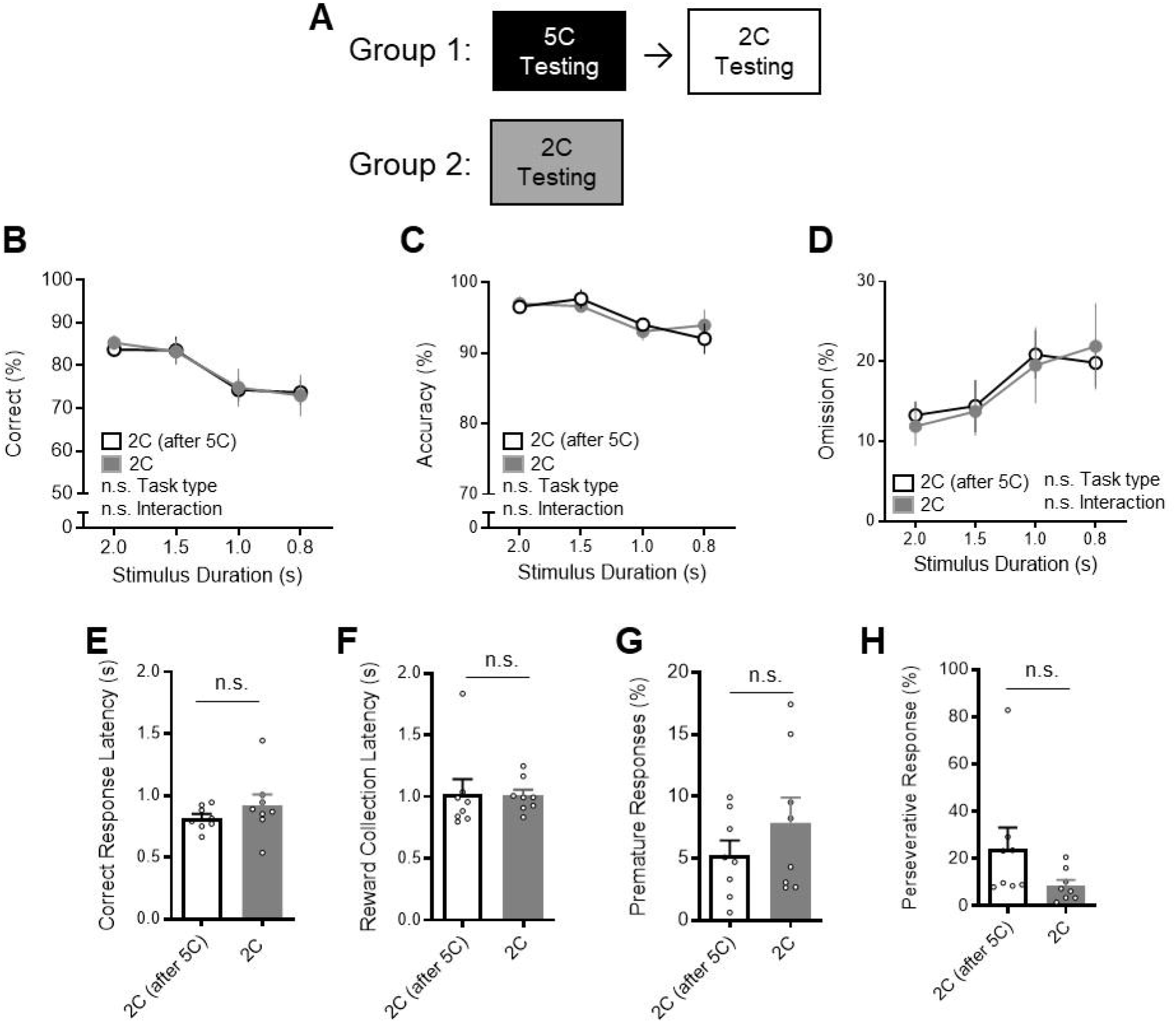
Prior 5CSRTT performance has no effect on 2CSRTT performance. **(A)** The impact of prior 5CSRTT testing on 2CSRTT testing was examined. Group 1 trained and tested on the 5CSRTT and then next performed the 2CSRTT testing. Group 2 trained and tested the 2CSRTT, without prior 5CSRTT testing. **(B)** There was no difference in correct % during the 2CSRTT between mice with prior 5CSRTT testing (n=8 mice) and mice without 5CSRTT testing (n=8 mice, two-way RM ANOVA, Main effect (task type): F_1,14_ =0.006283, *P*=0.9379, Main effect (stimulus duration): F_2.465.34.51_ =13, *P*<0.0001, Interaction: F_3,42_ =0.1039, *P*=0.9573). **(C)** There was no difference in accuracy % during the 2CSRTT between mice with prior 5CSRTT testing and mice without 5CSRTT testing (two-way RM ANOVA, Main effect (task type): F_1,14_ =0.004251, *P*=0.9489, Main effect (stimulus duration): F_3,42_ =5.225, *P*=0.0037, Interaction: F_3,42_ =0.5455, *P*=0.6539). **(D)** There was no difference in omission % during the 2CSRTT between mice with prior 5CSRTT testing and mice without 5CSRTT testing (two-way RM ANOVA, Main effect (task type): F_1,14_ =0.006206, *P*=0.9383, Main effect (stimulus duration): F_2.525,35.35_ = 8.264, *P*=0.0005, Interaction: F_3,42_ =0.3126, *P*=0.8162). **(E-H)** No difference in E) correct response latency (t_14_=1.049, *P*=0.3119), **(F**) reward collection latency (t_14_=0.08138, *P*=0.9363), **G**) premature responses (t_14_=1.097, *P*=0.2913), and **(H)** perseverative responses (t_14_=1.701, *P*=0.1111) was observed between the two groups. Error bars indicate mean□±□s.e.m. n.s. = non significant.

### 3.2 ACA_VIS_ projections, while required for 5CSRTT, are dispensable for 2CSRTT

While our previous studies have shown that ACA_VIS_ projection neurons are required for attentional behavior during 5CSRTT in mice (Norman et al., 2021a; Norman et al., 2021b), it is unknown whether the contribution of this subpopulation of frontal cortex projecting neurons are task demand dependent. To this end, we selectively suppressed ACA_VIS_ axon terminal activity in visual cortex using the inhibitory opsin halorhodopsin eNpHR3.0 at key time points during both the 5CSRTT and 2CSRTT in which the stimulus duration length was shuffled between 2s and 1s in a pseudorandomized order to determine its effect on task performance (**Fig.3A-B**). Data of the 5CSRTT have been analyzed previously (Norman et al., 2021b), but only the mice which underwent both 5CSRTT and 2CSRTT (6 out of 8 mice) were re-analyzed in this study. Viral spread and implant location were validated and reported in a previous study that used the mice to this study (Norman et al., 2021b). This study also showed that patch-clamp recording from ACAvis neurons in slices effectively suppressed spike activities without rebound excitation (Norman et al., 2021b).

**Figure 3.**
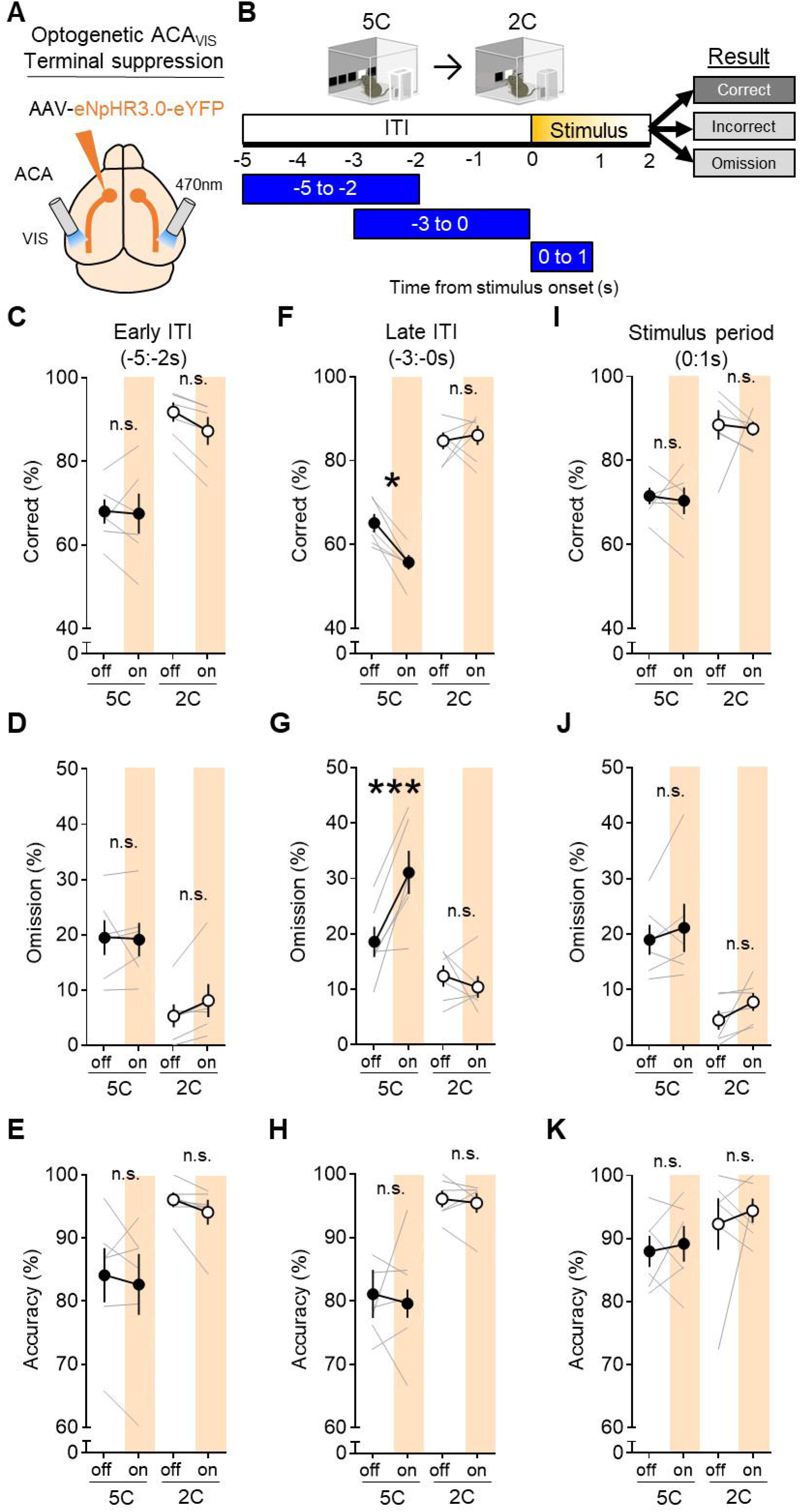
ACA_VIS_ projections, while required for 5CSRTT, are dispensable for 2CSRTT. **(A)** Viral strategy: To express light-gated chloride channel eNpHR3.0 in excitatory ACA neurons, eNpHR3.0 encoding AAV under the CaMKII promoter was injected in the ACA. An optic fiber was implanted at the VIS to optically suppress ACA_VIS_ projection terminals with temporal precision during the 5CSRTT. **(B)** Experimental timeline: Mice were first trained on the 5CSRTT, before viral injection. After allowing three weeks for maximal viral expression, mice underwent 5CSRTT testing. During 5CSRTT testing, with a fixed 5s ITI and pseudorandomized stimulus duration (2.0 or 1.0s), 470nm LED was illuminated during early ITI (−5:−2s), late ITI (−3:0s), or during period of stimulus presentation (0:1s) during 50% of trials (6 mice; 2,694 total trials). One timing was tested per day. The effect of optogenetic suppression during early ITI, late ITI, or stimulus period on behavioral performance (correct %, omission %, and accuracy %) at each light timing tested. The same mice were then baselined with 2CSRTT before undergoing the same optogenetic manipulation protocol with the 2CSRTT testing (6 mice; 2,844 total trials). **(C-E)** Continuous 470nm illumination during ITI −5:−2s had no effect on correct trials (%, general linear mixed model (GLMM), effect of light: *P* = 0.1010, effect of task: *P* < 0.0001, interaction: *P* = 0.1412), omissions (GLMM, effect of light *P* = 0.2873, effect of task: *P* < 0.0001, interaction: *P* = 0.2214), or accuracy (GLMM, effect of treatment (light on vs. off): *P* = 0.1796). **(F-H)**_Continuous 470nm illumination during ITI −3:0s had an effect on correct trials (GLMM, effect of light: *P* = 0.1274, effect of task: *P* < 0.0001, interaction: *P* = 0.0305) and omissions (GLMM, effect of light *P* = 0.0470, effect of task: *P* < 0.0001, interaction: *P* = 0.0008). Light disrupted correct trials during the 5C task (\**P* = 0.0114), but not during the 2C task (*P* = 0.9784). Light disrupted omissions during the 5C task (\*\*\**P* = 0.0001), but not during the 2C task (*P* = 0.8267). Continuous 470nm illumination during ITI −3:0s had no effect on accuracy (GLMM, effect of light: *P* = 0.6049, effect of task: *P* < 0.0001, interaction: *P* = 0.6068). **(I-K)** Continuous 470nm illumination during the stimulus period had no effect on correct trials (GLMM, effect of light: *P* = 0.4721, effect of task: *P* < 0.0001, interaction: *P* = 0.7455), omissions (GLMM, effect of light: *P* = 0.0242, effect of task: *P* < 0.0001, interaction: *P* = 0.1570, effect of light during 5C task *P*=0.6911, during 2C task *P*=0.9702), or accuracy (GLMM, effect of light: *P* = 0.2600, effect of task: *P* = 0.0009, interaction: *P* = 0.5844). Error bars indicate mean ± s.e.m. n.s. = non significant. * indicates *P*<0.05, *** indicates *P*<0.001.

Correct response rate provides a broad and general index of attention performance and reflects what is captured by both accuracy and omission, two readouts that have been used to assess some aspects of attention in previous studies using 5CSRTT (Carli et al., 1983). We found that suppressing ACA_VIS_ projections during the 3s immediately before stimulus presentation (−3:0s) impaired performance during the 5CSRTT, reducing correct trials (**Fig.3F** general linear mixed model (GLMM) effect of light during 5C task *P* = 0.0114) by increasing omission (**Fig.3G** effect of light during 5C task *P* = 0.0001), however, no effect was observed during the 2CSRTT (**Fig.3FG** GLMM, effect of light during 2C task: correct *P* = 0.9784, omission *P* = 0.8267). We did not observe effect of light illumination on accuracy (**Fig.3H** GLMM, effect of light *P* = 0.6049). These findings indicate that it is mainly the omission increase that contributed to the correct rate reduction by the optogenetic suppression of ACAvis projections during 5CSRTT.

Performance of either task was not affected by adjusting the LED illumination time to earlier in the ITI (−5:−2s) (**Fig.3C-E**: GLMM, effect of light: correct trials %*P* = 0.1010, omissions %*P* = 0.2873, accuracy %*P* = 0.1796) or during stimulus presentation (0:1s) (**Fig.3 I-K**: GLMM, correct trials %: effect of light *P* = 0.4721, omission: effect of light during 5C task P=0.6911, during 2C task P=0.9702, accuracy%: effect of light *P* = 0.2600). Of note, our recent study showed that LED illumination over VIS to control mice expressing eYFP in ACA induced no behavioral effects during 5CSRTT (Norman et al., 2021b), suggesting that behavioral changes depend on halorhodopsin expression and are unlikely to be due to other factors such as light induced heat change. It should be though noted that eYFP control experiments were not performed during 2CSRTT. ACA_VIS_ terminal inhibition during any time point examined had no effect on reward collection latency (**Fig. S2A**: t test, ITI −5:−2s *P*=0.3846, ITI −3:0s *P*=0.7773, stimulus period *P*=0.9072), correct response latency (**Fig. S2B**: t test, ITI −5:−2s *P*=0.7012, ITI −3:0s *P*=0.2568, during stimulus period *P*=0.7620), premature responses (**Fig. S2C**: t test, ITI −5:−2s *P*=0.1002), or perseverative responses (**Fig. S2D**: t test, ITI −5:−2s *P*=0.5593, ITI −3:0s *P*=0.5554, stimulus period *P*=0.2756) during the 2CSRTT.

Taken together, these data suggests that ACA_VIS_ projections are causally important for attentional task performance in the period just before stimulus presentation during 5CSRTT, but become dispensable once the task switch to 2CSRTT, suggesting that ACA_VIS_ projections are necessary only when task demand is high.

## 4. Discussion

In this study, we first characterized how attentional behavior outcomes depend on the number of locations for mice to scan and touch for successful performance, and found that the 2CSRTT is less task demanding than the 5CSRTT. We then used circuit-specific optogenetic suppression to show that frontal-sensory ACA_VIS_ projection neurons are only crucial for top-down control of attention behavior in male mice during high task demand conditions. These findings support a model that the frontal-sensory ACA_VIS_ projection regulates visual attention behavior during specific high task demand conditions. Previous studies in mice revealed a causal role of a specific sub-population ACA neurons projecting to the visual cortex in contributing to attentional behavior (Norman et al., 2021a; Norman et al., 2021b), similar to other ACA neurons (Kim et al., 2016; White et al., 2018). However, it remained unclear whether this mechanism was dependent on task demand. Our study provides further insight into defining the role of the specific factor of task demand has in recruiting particular subtypes of neurons.

There are some important limitations with the current study which warrant future investigations. While we showed that prior 5CSRTT training did not affect 2CSRTT performance, one limitation of the current study is that all mice in optogenetic experiments performed the 5CSRTT prior to the 2CSRTT. Future studies should randomize the order in which mice are trained on each of the task in order to control for the contribution of ACA_VIS_ neurons to attentional behavior. It would also informative to investigate the activity of ACAvis neurons during 2CSRTT and 5CSRTT by combining *in vivo* electrophysiology to examine to what extent ACAvis neuron activity reflects task demand. It should be also noted that this study manipulated a number of choice locations between tasks, but not other parameters which can also impact task demand such as the duration of ITI or visual stimulus. Future studies are warranted to examine to what extent task demand-dependent aspect of ACA_VIS_ function is generalizable.

Our study provides insight into the neural mechanisms underlying cognitive behavior deficits in psychiatric disease. In low attentional load conditions, attention function while ignoring visual distractors does not differ between patients with schizophrenia and healthy controls but those with schizophrenia performed worse during high load conditions, suggesting that attentional deficits only emerge under more challenging tasks (Ducato et al., 2008). Our study may ultimately inspire consideration of task demand when designing and interpreting the results of interventions specifically targeting top-down frontal-sensory circuits to improve attention.

## 6. Author Contributions

KJN and HM designed experiments. KJN and HM wrote the manuscript with contributions from all co-authors. KJN performed behavioral and optogenetics experiments and analyzed data with assistance from JB, HK, and TN. JB, PM, CC, and KC performed behavioral training. HM supervised the research.

## 8. Acknowledgments

We thank Drs. Mark Baxter and Peter Rudebeck for feedback on behavioral and statistical analysis. We also thank members of the Morishita lab for helpful feedback.

