## Supplemental figure 1 and 2 for "Frontal-sensory cortical projections become dispensable for attentional performance upon a reduction of task demand in mice"

### Supplemental Material

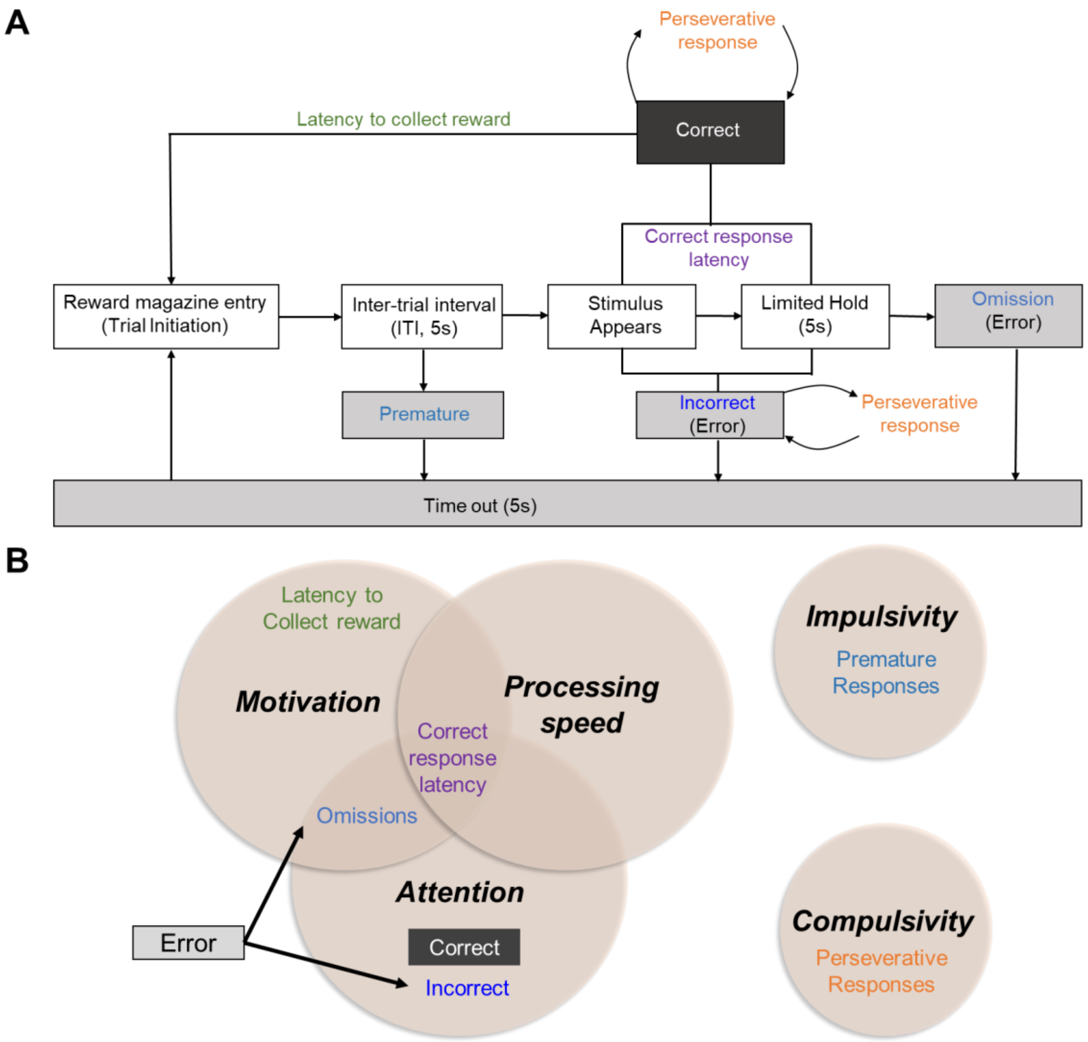


**Figure S1. Overview of 5CSRTT and 2CSRTT protocol. A)** Schematic of 5CSRTT testing. A trial begins when the reward magazine is entered. Upon exiting the reward magazine, a 5 second intertrial interval (ITI) begins. After the ITI, a white square stimulus will randomly appear at 1 of 5 touchscreens for either 2.0, 1.5, 1.0, or 0.8 s (2.0 or 1.0 s for optogenetic experiments). If one of the touchscreens are touched during the ITI, it's considered a premature response and a 5s timeout is initiated. If the correct touchscreen is touched when the stimulus is present or during the following 5s (limited hold) a milk reward is dispensed in the reward magazine. However, if the incorrect location is touched, a 5s timeout is initiated. If none of the locations are touched, a 5s timeout is initiated (omission). If the touchscreen is touched again after either a correct or incorrect response, it's considered a perseverative response. Correct response latency refers to the time from stimulus presentation to correct response. Latency to collect reward refers to the time from correct response to reward tray entrance. **B)** Overview of cognitive functions that contribute to various actions during 5CSRTT performance. While there is some overlap in measurements used to assess attention during the 5CSRTT, each measure has a specific purpose and represents various levels and specificity of attention capacity. Correct response rate is a broad and overall index of attention capacity which reflects what is captured by both accuracy and omission, two readouts which are used to assess attention level in previous studies using

5CSRTT. Placing ‘Omissions’ at the intersection of attention and motivation in the Venn diagram emphasizes that omission probability is sensitive to both. However, another readout of the 5CSRTT, the latency to collect reward following a correct response, serves as an indicator of motivation to assess if omission change reflects attention or motivation level. Premature responses that occur prior to the stimulus appearing on the touchscreen are excluded from error trials in our analysis because premature trials are rather interpreted as errors of impulsivity as the mice are unable to wait or inhibit a response during the 5s ITI (Robbins, 2002).

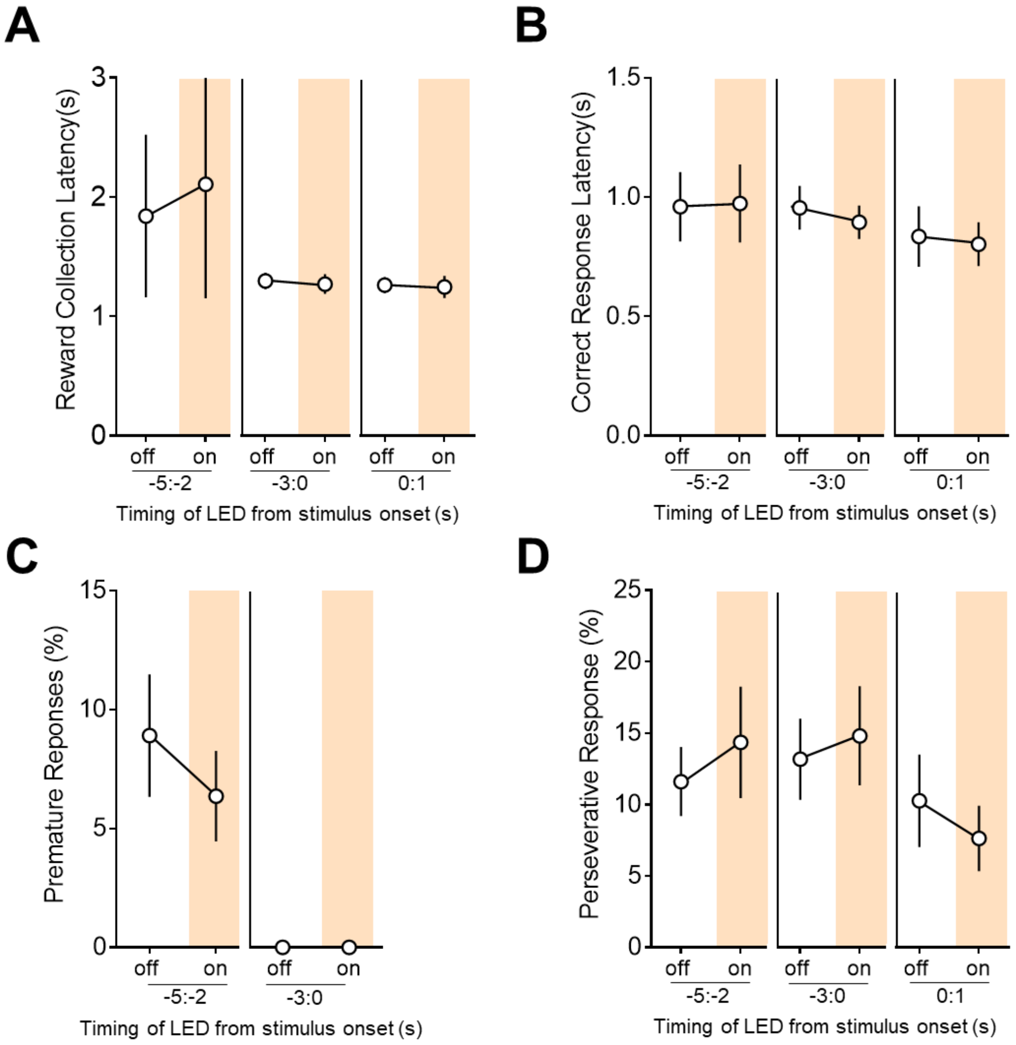


**Figure S2. Optogenetic suppression of ACA$_{VIS}$ projections during 2CSRTT**. **(A)** Reward collection latency was not affected by ACA$_{VIS}$ suppression during ITI -5:-2s ($t_5$=0.9526, $P$=0.3846, n=6 mice), ITI -3:0s ($t_5$=0.2985, $P$=0.7773, n=6 mice) or stimulus period ($t_5$=0.1226, $P$=0.9072, n=6 mice) compared to light off trials. **(B)** Correct response latency was not affected by ACA$_{VIS}$ suppression during ITI -5:-2s ($t_5$=0.4064, $P$=0.7012, n=6 mice), ITI -3:0s ($t_5$=1.280, $P$=0.2568, n=6 mice) or during stimulus period ($t_5$=0.3198, $P$=0.7620, n=6 mice) compared to light off trials. **(C)** Premature responses were not affected by ACA$_{VIS}$ suppression during ITI -5:-2s ($t_5$=2.014, $P$=0.1002, n=6 mice) or ITI -3:0s compared to light off trials. The effect of suppression during stimulus period on premature responses was not analyzed, because premature responses occur before the stimulus appears. **(D)** Perseverative responses were not affected by ACA$_{VIS}$ suppression during -5:-2s ($t_5$=0.6252, $P$=0.5593, n=6 mice), ITI -3:0s ($t_5$=0.6316, $P$=0.5554, n=6 mice) or stimulus period ($t_5$=1.224, $P$=0.2756, n=6 mice) compared to light off trials.
